# Repositioning linifanib as a potent anti-necroptosis agent for sepsis

**DOI:** 10.1101/2022.03.24.485557

**Authors:** Kai Yang, Min Li, Liang Yu, Xiaoyan He

## Abstract

**Background:** Sepsis is a systemic inflammatory syndrome (SIRS) caused by acute microbial infection with high mortality rate. The role of tumour necrosis factor α (TNF-α)-induced necroptosis in promoting the pathophysiology of sepsis has been identified. Effective prevention of necroptosis is expected to improve the prognosis of sepsis patients.

**Methods:** We conducted bioinformatics prediction of candidate drugs by analyzing differentially expressed genes of sepsis patients extracted from GEO database, combining library of integrated network-based cellular signatures (LINCS) L1000 perturbation database. Biological experiments based on TNF-α-induced necroptosis in cellular and mouse model were performed to verify the protection of candidate drugs from SIRS. Cell viability was measured by CellTiter-Glo luminescent ATP assay. Effects of linifanib on necroptosis were investigated by western blotting, immunoprecipitation, and in vitro RIPK1 kinase assay. Survival curve analysis of SIRS mice treated by linifanib was performed.

**Results:** A total of 16 candidate drugs was screened out through bioinformatics analysis. Our experiments demonstrated that linifanib effectively protected cells from necroptosis and rescued the death of SIRS mice from shock induced by TNF-α. In vitro, linifanib directly suppressed RIPK1 kinase activity. In vivo, linifanib effectively reduced the overexpressed level of IL-6, a good marker of severity during severe sepsis, in the lung of SIRS mice.

**Conclusion:** We provide preclinical evidence for the potential clinical utility of linifanib in sepsis. Study of drug repositioning using bioinformatical predictions combined with experimental validations provides novel strategies for the development of sepsis drug.

## Introduction

Sepsis is a systemic inflammatory syndrome (SIRS) caused by acute microbial infection, at 30-50% mortality rate though antibiotics are widely applied for the treatment (Choi et al. 2020; Martínez-García et al.; Iskander et al. 2013). Given the high morbidity and mortality of sepsis and septic shock, the Surviving Sepsis Campaign was initiated globally to help improve the treatment of sepsis and septic shock(J.-S. et al. 2011). Consequently, a great effort is needed to find new, and more effective therapeutic agents for sepsis/septic shock(Chiao et al. 2013). In this study, we sought to employ a systematic drug repositioning(Katsila et al. 2016; Lavecchia and Giovanni 2013) bioinformatics approach to identify novel FDA-approved candidate drugs to treat sepsis (Pushpakom et al. 2019). We proposed a new approach to identify therapeutic drugs for sepsis from existing small molecular compounds, by investigating the difference of functional pathways and the gene expression profile of blood nucleated cells in patients with sepsis and normal control group. This method comprehensively considers the genome expression profile in the state of sepsis, and finds the drugs to reverse the changes of gene expression through pattern matching.

Injury of multiple organs induced by cytokine storm and SIRS underlies sepsis progression with septic shock. The role of Tumour necrosis factor α (TNF-α) - induced necroptosis in promoting the pathophysiology of sepsis has been identified. Necroptosis induced by TNF-α initiates downstream signaling cascades driving the production of a wide array of pro-inflammatory cytokines. Necroptosis mediated SIRS and associated pro-flammatory cytokine storms are increasingly recognized as an important pathogenic mechanism of sepsis(Duprez et al. 2011; Lei et al. 2014). Blocking TNF-α-induced necroptosis might be useful in mitigating the cytokine storm and preventing severe sepsis clinically. Thus, effective prevention of necroptosis is expected to improve the prognosis of sepsis patients(Wynn et al. 2008). However, no mature inhibitors of necroptosis are currently in clinical use(Moerke et al. 2019). Therefore, the role of candidate drugs from our previous bioinformatics analysis was verified by the TNF-induced necroptosis cellular model in vitro and the TNF-α induced SIRS mouse model in vivo in our study.

Our study of drug repositioning using bioinformatical predictions combined with experimental validations is expected to provide novel strategies for the development of drug for sepsis.

## Results

### Initial identification of candidate drugs to treat sepsis by disease-drug associations

We found three sepsis-related datasets from GEO database. Bioinformatics analysis was carried out on these datasets to determine the differentially expressed genes in sepsis compared with normal state. We identified signaling pathways related to the pathogenesis of sepsis from the literature. Finally, according to the gene expression data in the sepsis-related pathways and the gene expression profile of human cell lines induced by small molecular compounds in L1000 library, a pattern matching method is designed based on Kolmogorov-Smirnov test. For each sepsis-related dataset, the treatment scores of all drugs in L1000 library were calculated, and the drugs were sorted. The top 60 drugs in the three datasets were verified in the CTD database, and the drugs unrelated to sepsis or without information in the database were selected as candidate drugs for sepsis. We identified 16 drugs with potential therapeutic effects on sepsis. The overall flow chart is shown in Figure 1.

**Figure1.**
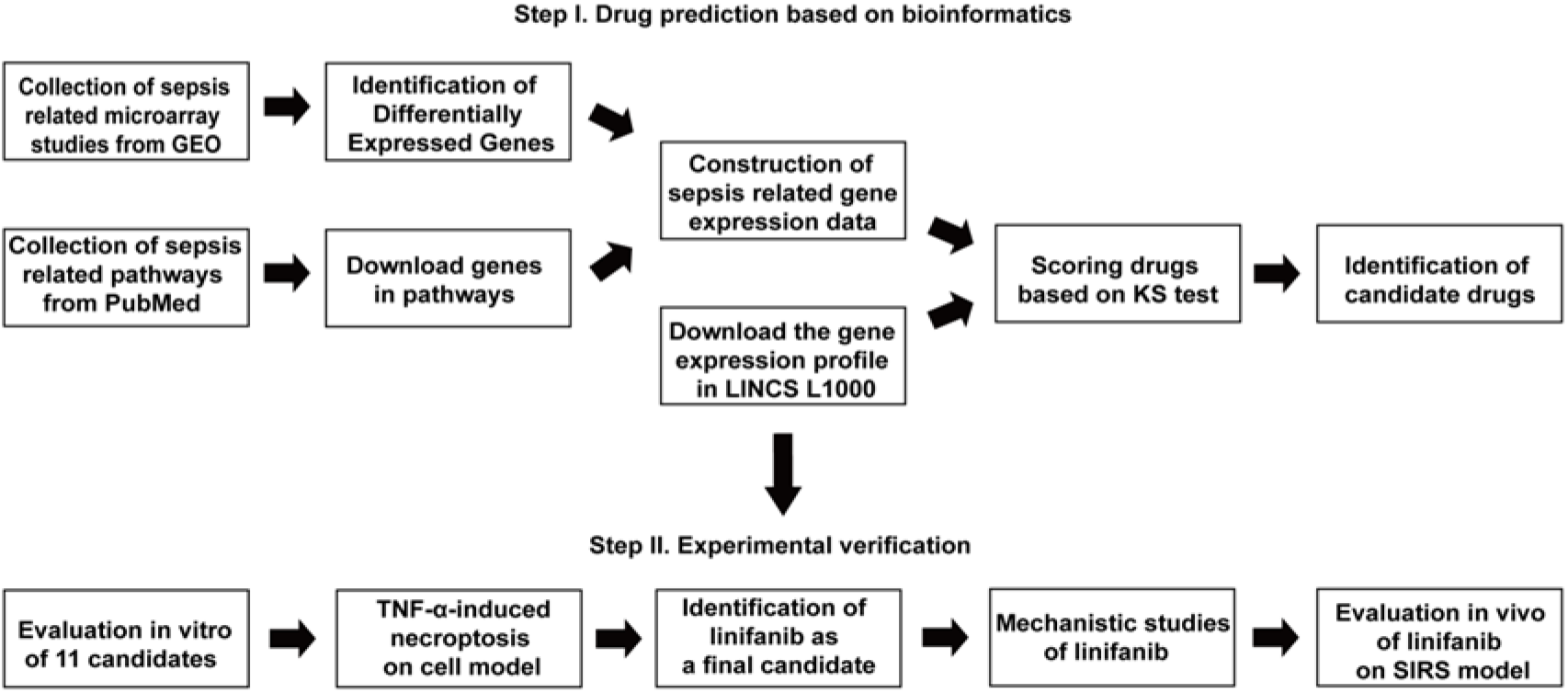
Experimental design. For the virtual screening for sepsis treatment, sepsis-related signaling pathways and datasets were found from public database. DEGs were extracted using sepsis-related microarrays from the GEO. Sepsis-related gene expression data and gene expression profiles in the L1000 database were used to calculate therapeutic scores to obtain candidate compounds. In the next step of experimental validation, FDA-approved drug linifanib was shown to have therapeutic effects in cellular and animal models of necroptosis.

### Identification of gene expression profiles of sepsis

We obtained the gene expression profiles of sepsis patients and healthy subjects from GSE69528, GSE46955 and GSE54514 datasets and performed differential expression analysis. We set P value < 0.05 and identified 7771, 6238, 5564 DEGs from the GSE69528, GSE46955 and GSE54514 datasets. We set |log2FC| > 0.7, and identified the DEGs that were significantly up-regulated or down-regulated in three datasets, and their overlapping DEGs are shown in Figure 2A. The volcanic plots are shown in Figure 2B. The heatmap of the TOP 100 DEGs in each dataset are shown in Figure 2C.

**Figure 2.**
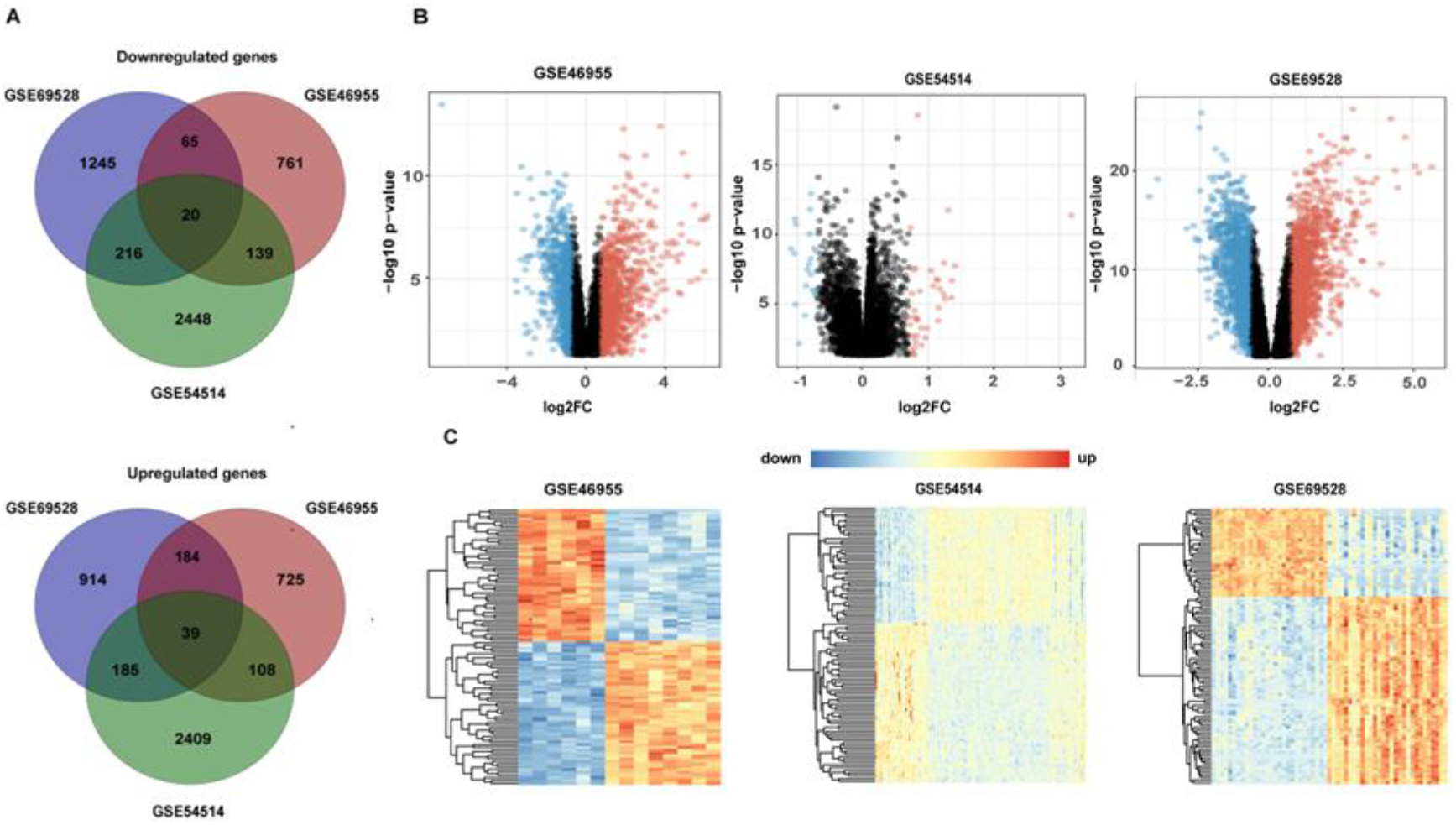
Differential expression analysis of GSE69528, GSE46955 and GSE54514 datasets. (A) Venn diagram shows the overlapping DEGs in the three datasets. (B) Volcano plot shows significantly changed genes in the three datasets. Red and blue plots represent up- and downregulated genes, respectively (|log2FC| > 0.7 and P value < 0.05). Black plots represent the genes with no significant difference. (C) Heatmap of the top 100 DEGs in three datasets. Each row represents a DEG, each column represents a sample. The color ratio indicates the relative level of DEG expression: blue, lower than the reference expression, red, higher than the reference expression.

### Detection of sepsis pathway signatures

This study searched the sepsis-related literature from the biomedical database, and identified 17 signaling pathways related to the pathogenesis of sepsis or important for the prognosis of sepsis patients, as shown in Table 2. These pathways may contain key pathogenic genes of sepsis, and the expression of these genes is different in patients with sepsis and healthy subjects. Therefore, it is necessary to focus on the gene sets involved in these signaling pathways and take them as the feature pathways of sepsis.

**Table 1.**
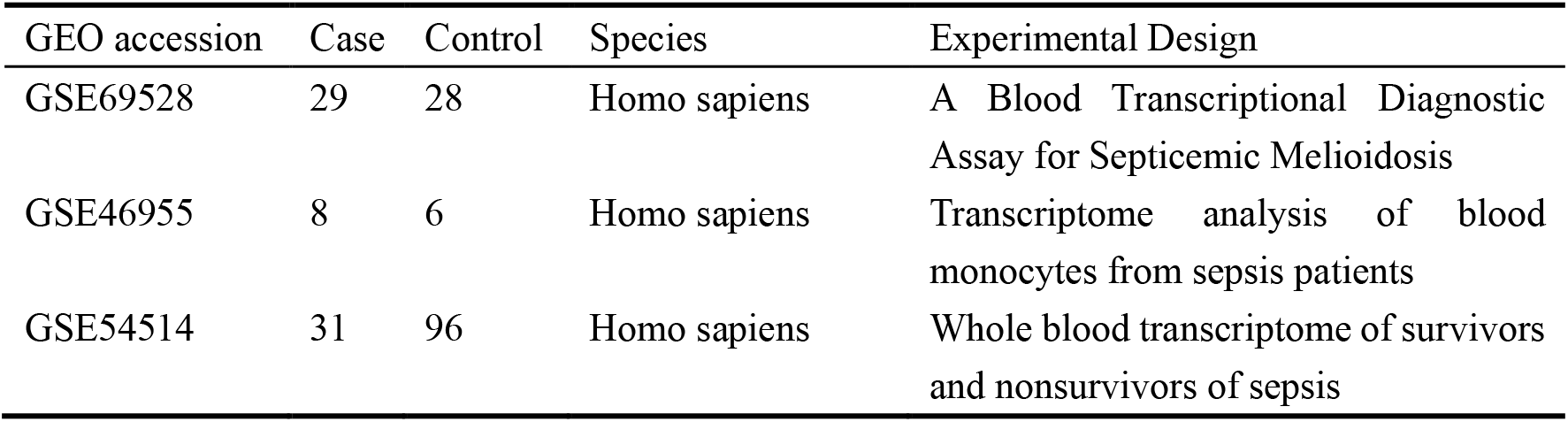
Description of GEO datasets used in this study.

**Table 2.** The mean value of logFC of DEGs in signaling pathways related to sepsis.

| Pathway | Mean value of logFC |
| --- | --- |
| TNF signaling pathway | 0.513 |
| Cytokine-cytokine receptor interaction | 0.409 |
| NOD-like receptor signaling pathway | 0.377 |
| Toll Like Receptor 4 (TLR4) Cascade | 0.319 |
| NF-kappa B signaling pathway | 0.318 |
| JAK-STAT signaling pathway | 0.279 |
| C-type lectin receptor(CLR) signaling pathway | 0.270 |
| Necroptosis | 0.213 |
| PD-L1 expression and PD-1 checkpoint pathway in cancer | 0.160 |
| The NLRP3 inflammasome | 0.134 |
| PI3K-Akt signaling pathway | 0.104 |
| AGE-RAGE signaling pathway in diabetic complications | 0.103 |
| MAPK signaling pathway | 0.093 |
| STING mediated induction of host immune responses | 0.063 |
| T cell receptor signaling pathway | 0.017 |
| mTOR signaling pathway | 0.003 |
| PPAR signaling pathway | -0.069 |

We integrated the logFC of DEGs in the three datasets with the genes in each pathway, and calculated the average value of logFC of all genes as the result of up-regulation or down-regulation of each pathway. Finally, we calculate the average value of logFC of the pathways corresponding to the three datasets, as shown in Table 2. We can see that the upregulation of TNF signaling pathway is the highest.

### Selection of potential sepsis drugs through CTD benchmark

We used the gene expression information of three datasets to calculate the therapeutic score, so we got three drug lists. First, these drugs were validated in a public database. The Comparative Toxicogenomics Database(CTD) provides information on the relationship between genes, compounds and diseases(Allan et al. 2018). In this study, we used CTD to query the association between compounds and sepsis. The drug lists of three datasets GSE46955, GSE69528 and GSE54514 were sorted, and the top 60 drugs were selected and verified in CTD. Drugs were divided into four categories: associated with sepsis, no compounds in CTD, no disease data with compounds, and not associated with sepsis. They are all candidate drugs for potential treatment of sepsis except drugs related to sepsis. The distribution of top 60 drugs of the three datasets is shown in Table 3.

**Table 3.**
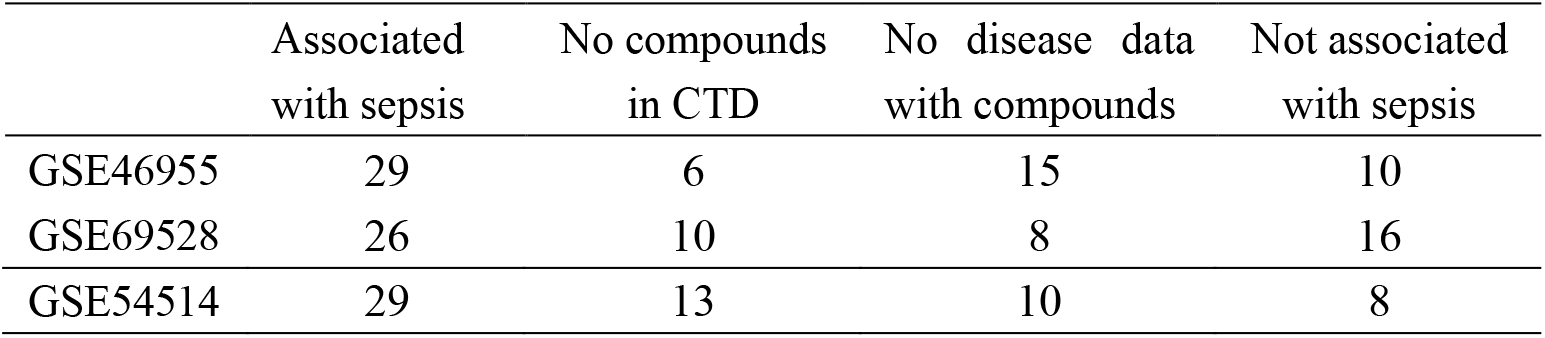
Distribution of the top 60 drugs of three datasets.

There are 16 candidate drugs included in two or three drug lists: Y-39983, CHIR-99021, WH-4-025, brivanib, XMD-1150, CGP-60474, saracatinib, enzastaurin, withaferin-a, AT-7519, linifanib, asenapine, nintedanib, AS-601245, GSK-1059615 and OSI-027. Three of them have previously been reported anti-sepsis effects in the literature: CGP-60474(Han et al. 2018), AT-7519(Dorward et al. 2017), and Y-3998(Tokushige et al. 2011) and 11 drugs have not yet been reported while no information for 2 drugs (WH-4-025, XMD-1150) is available. The overlap of candidate drugs is shown in Figure 3.

**Figure3.**
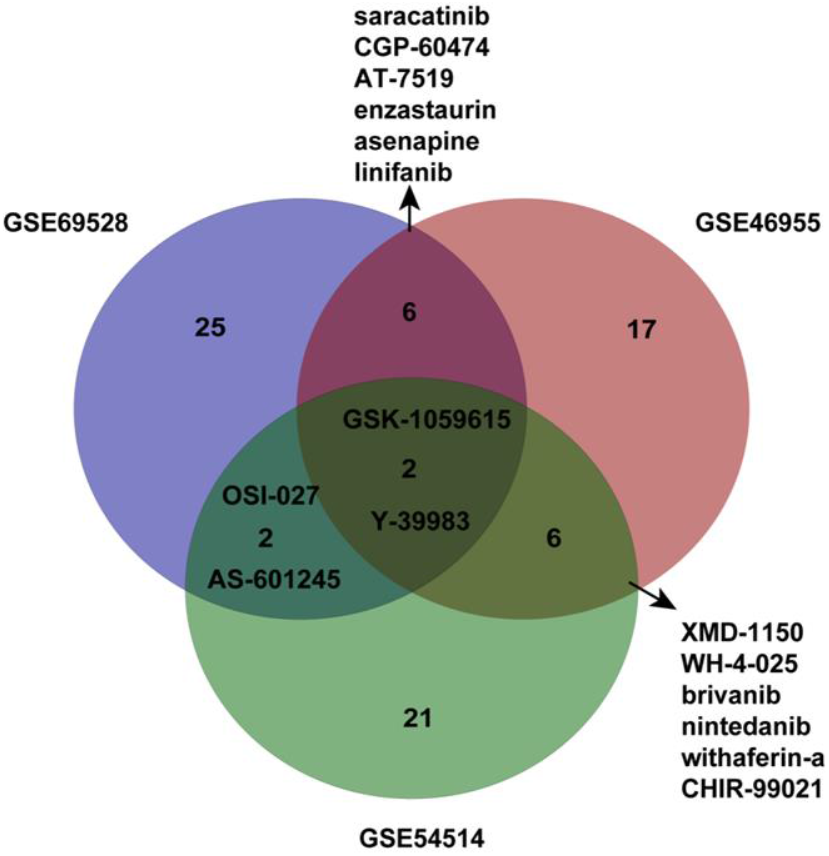
Venn diagram shows candidate drugs for sepsis obtained from the three datasets. 16 drugs appeared in at least two results.

### Experimental validation of potentially anti-sepsis drugs via in vitro cellular assay

Necroptosis plays a significant role in the pathophysiology of SIRS and sepsis. RIPK1 regulates RIPK3-MLKL-driven systemic inflammation and RIPK1 kinase inhibitors show promise in alleviating or preventing a SIRS response. Our study found that the expression of RIPK1, RIPK3 and MLKL in peripheral blood nucleated cells of sepsis patients was significantly higher than that of healthy controls(Figure 4A). These results provided a useful reservoir for further experimental verification.To further explore the potential novel therapy target of our bioinformatics predictions, we performed a phenotypic screen for drugs analysis as inhibitors of TNF-α -induced necroptosis in FADD-deficient Jurkat cells. FADD - deficient Jurkat cells were pretreated with or without these compounds (10 μM, 30 min) and then stimulated with 50ng/ml TNF-α for 24 h or left untreated(Lawrence and Chow 2005). Nec-1 was used as positive control to monitor RIPK1 kinase dependency in the induced necroptosis.(Moriwaki and Chan 2014). Cell viability was determined using the ATP-based CellTiter-Glo® Luminescent Cell Viability Assay (Promega). Measurements were normalized to those in untreated control cells (100%) (Figure 4B). Our study demonstrated that 1 (linifanib) of 16 candidate drugs protected effectively cells from necroptosis (Figure4 C-E).

**Figure 4.**
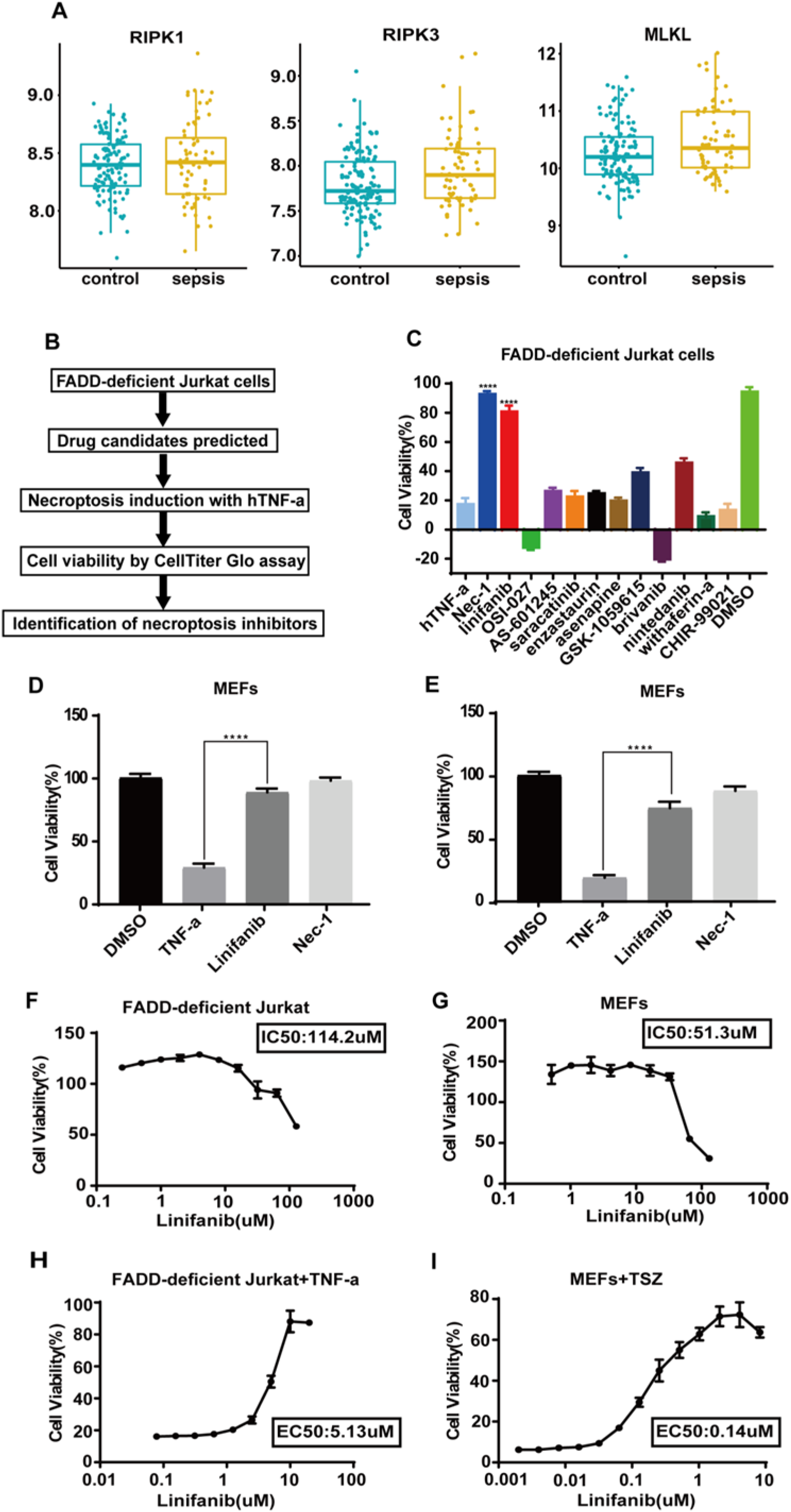
Experimental validation of potentially sepsis-related drugs via in vitro cellular assay. (A) Boxplots of the essential genes(RIPK1,RIPK3 and MLKL) in necroptosis pathway in sepsis three datasets. (B)Schematic overview of the drug screen workflow. (C) Identification of necroptosis inhibitor by using candidate drug predict with LINCS and CTD databases for the treatment of sepsis based on bioinformatics. FADD-deficient Jurkat cells were pretreated with each compound (10 μM) for 30 min and then stimulated with human TNF-α (50 ng/ml) overnight to induce necroptosis. (DMSO= 100). Cell viability of FADD-deficient Jurkat cells (D) andMEFs cells (E) which were pretreated with linifanib (10 μM) for 30 min in the presence or absence of 25 μM Nec-1 and then stimulated with 50 ng/ml human TNF-α or TSZ:TNF - α (25 ng/ml) plus Smac mimetic (200 nM), and z - VAD - fmk (20 μM) for 24 hr. Dose-response protection of linifanib in (F) FADD-deficient Jurkat cells and (G) MEFs cells were incubated for 24 hrs with linifanib (0.05 μM-100 μM). Cell viability was determined in dose-dependent protection of linifanib in (H)FADD-deficient Jurkat cells treated with various concentrations of linifanib (0.01 μM-20 μM) followed by stimulation with human TNF-α (50 ng/ml) for 24hrs, while (I)MEFs cells were pre-incubated with linifanib (concentrations as indicated) for 30 minutes with 25 ng/ml mTNF-α (T) together with 200 nM Smac164 (S) and 20 μM caspase inhibitor z-VAD (Z) for 24 hr. Cell viability was assessed using a luminescence-based readout for ATP (CellTiter Glo) throughout. Data represent mean value±S.D. from three independent experiment and normalized to untreated control. ****p < 0.0001, significant effect of linifanib treatment.

We further performed dose–response curves to quantitatively assess the inhibitory potency of linifanib. Drug toxicity of linifanib was assessed by determining the half maximal inhibitory concentration (IC50), which was 114.2 uM in FADD-deficient Jurkat cells(Figure 4F) and 51.3 uM in MEFs cells(Figure 4G). In addition, the half maximal effective concentration (EC50) for inhibiting necroptosis in this setting was measured to be 5.13μM in human FADD-deficient Jurkat cells (Figure 4H), 0.14 uM in MEFs cells (Figure 4I). Highlighting the window of opportunity for necroptosis inhibition. Taken together, these data demonstrated that linifanib efficiently blocked TNF-α-induced necroptosis in both human and murine cells, was a potent necroptosis inhibitor.

### Linifanib blocks necrosome formation by inhibiting the phosphorylation of RIPK1

We found that the protective effect of linifanib in TNF-α -induced necroptosis was as potent as that of the well-established but not clinically applicable RIPK1 inhibitor Nec-1(Riebeling et al. 2020). We next evaluated whether the inhibitory mechanism of linifanib on necroptosis requires the involvement of RIPK1. FADD-deficient Jurkat cells were pretreated with 4 μ M linifanib or 25 μ M Nec-1 for 30min followed by treatment with or without TNF-α for indicate time points. Linifanib selectively inhibited the phosphorylation RIPK1 and MLKL when necroptosis was induced by TNF-α in FADD-deficient Jurkat cells(Figure 5A). Similar results were also obtained in murine MEFs cells (Figure 5B). Thus, these results indicate that linifanib inhibits TNF-α - induced phosphorylation of RIPK1 and MLKL.

**Figure 5.**
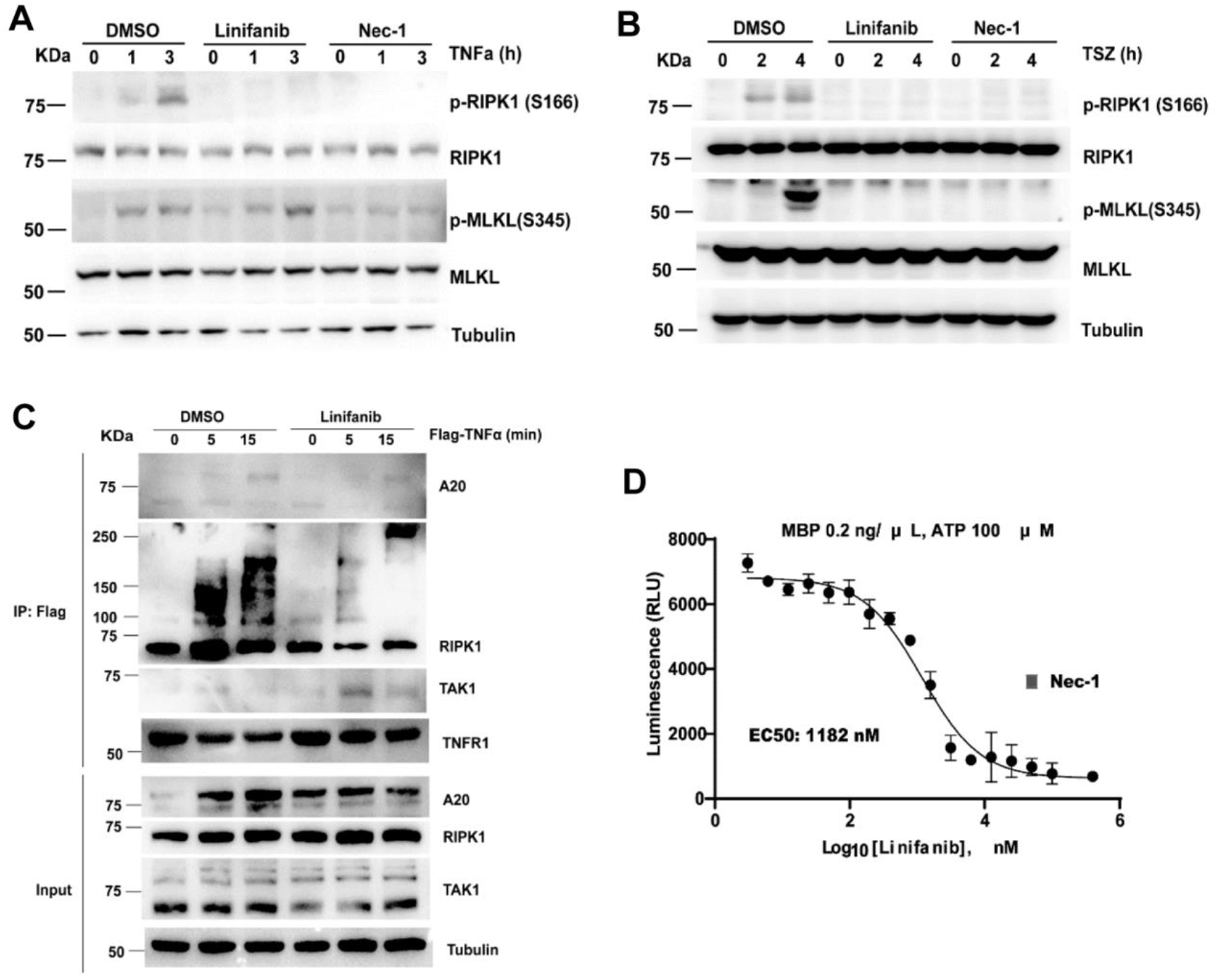
Linifanib directly binds to RIPK1 and inhibits the kinase activity. (A) FADD-deficient Jurkat cells were pretreated with linifanib (10 μM), Nec-1 (25 μM) or vehicle(0.2%DMSO) for 30 min, then incubated by hTNF-α (50 ng/mL) for indicated time. The levels of p-RIPK1^S166^ and p-MLKL^S345^ were determined by western blotting. (B) MEFs were pretreated with linifanib (5 μM), Nec-1 (25 μM) or vehicle for 30 min, then incubated with SM164 (200 nM) and z-VAD-fmk (20 μM) followed by mTNF-α (25 ng/mL) for indicated time. The levels of p-RIPK1^S166^ and p-MLKL^S345^ were determined by western blotting. (C) MEFs were pretreated with linifanib (0.31μM) or vehicle for 30 min and stimulated with Flag-mTNF-α (100 ng/ml) for indicated time points, and the complex I was pulled down using anti-Flag beads and analyzed by western blotting with the indicated antibodies. (D) Quantification of ADP - Glo kinase assays performed with recombinant hRIPK1 in the presence of increasing concentrations of linifanib. RIPK1 inhibitor Nec-1 was used as a positive control. Data represent mean value ± S.D. of two independent experiments performed in triplicates and normalized to untreated control.

TNF-α can rapidly activate TNFR1 upon binding, thereby inducing the recruitment of RIPK1, TAK1 to TNFR1 to form a TNFR1 signaling complex (TNF-RSC or complex I)(Song et al. 2021; Fan et al. 2011; Lei et al. 2014). Analysis of complex I formation influenced by linifanib was performed immunoprecipitated (IP) after stimulation with anti-TNFR1 antibody FLAG-tagged TNF-α using mouse primary fibroblastic cells(Riebeling et al. 2020; Nakazawa et al. 2016). We found that the recruitment of RIPK1 into the TNF-RSC was largely blocked in TNF-α stimulated-MEFs by linifanib. In addition, the recruitment of TAK1 was increased in linifanib-pretreated MEFs(Figure 5C).

As linifanib suppressed active RIPK1 kinase-mediated necroptosis and SIRS in response to TNF-α. We then conducted an in vitro kinase assay to further test the potency of linifanib directly inhibit RIPK1. Quantification of ADP-Glo kinase assay showed that linifanib suppressed recombinant hRIPK1(Rojas-Rivera et al. 2017) kinase activity (IC 50 = 1182 nM) (Figure 5D). These data demonstrated molecular mechanisms by which linifanib protect cells from TNF-induced necroptosis. As demonstrated by an in vitro kinase assay, linifanib directly suppressed the kinase activity of RIPK1. Our foundings indicated that linifanib is as effective as the well established RIPK1 inhibitor Nec-1 in preventing necroptosis and is a RIPK1 kinase inhibitor.

### Linifanib protects mice from TNF-α-induced SIRS

To explore whether linifanib protects against inflammation in vivo, we tested it in the TNF-induced SIRS model(Duprez et al. 2011). Linifanib (50mg/kg) or Nec-1 (30mg/kg), a positive control as an effective inhibitor of necroptosis, given by intragastric gavage 30 min before i.v. injection of mTNF-α, protected mice from hypothermia and death (Figure 6 A,B). Since interleukin 6 (IL-6) level in vivo correlated well with TNF alpha serum level and the mortality rate of patients with septic shock(Hou et al. 2015; Dama et al. 1992; Shahkar et al. 2011), we detected the expression level in the lung of SIRS mice. Our data showed that the SIRS mice showed significantly higher levels of IL-6 expression in the lung tissue at 6 hours after the mice received the TNFA injection in the tail vein, compared with normal control mice. And the overexpressed IL-6 of SIRS mice was significantly suppressed by pretreatment with linifanib. Thus, these results demonstrate that linifanib protects against TNF-induced SIRS in vivo.

**Figure 6.**
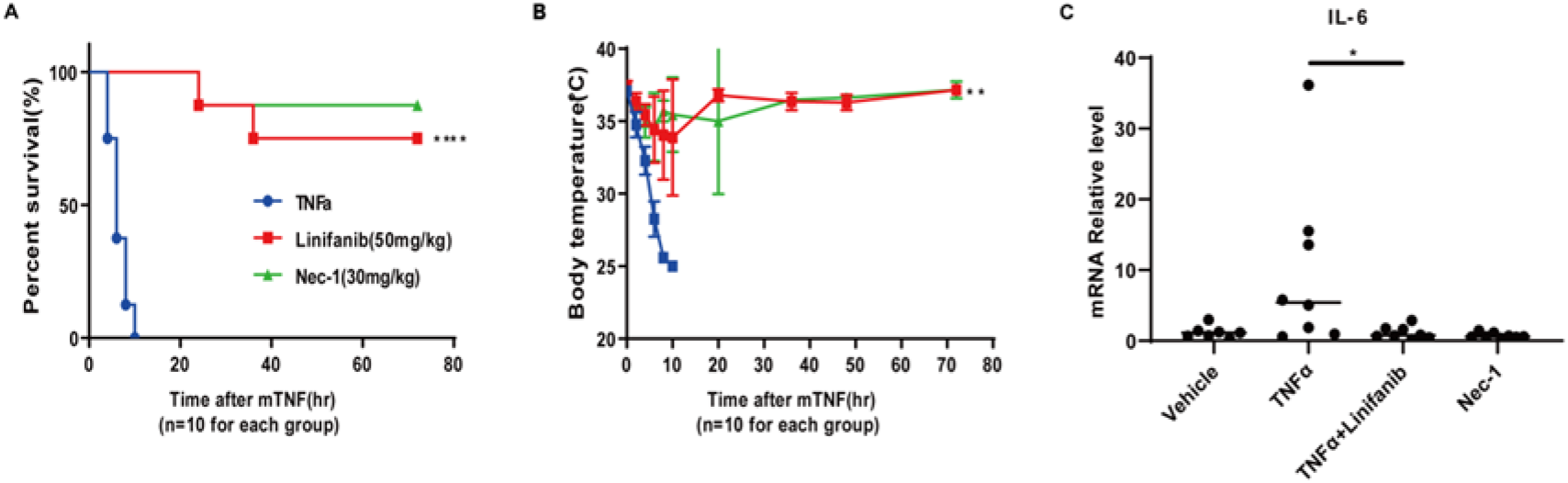
Linifanib protects against SIRS induced by TNF-α. 10-week-old male C57BL/6 J mice were pretreated with or without linifanib (50 mg·kg-1 gavage) or Nec-1 (30 mg·kg-1 gavage) for 30 minutes, and then SIRS was induced with mTNFα(5 μg/mice i.v). (A)The survival curve and (B)the body temperature changes (means ± SEM) of the vehicle control and linifanib - treated mice or Nec-1-treated mice (n = 10 for each group) are shown. (C) After SIRS induction for 6 hr, the IL-6 levels of lung tissues were determined by real-time PCR. *P < 0.05, **P < 0.01, ****P < 0.0001, significantly different from vehicle control group.

## Discussion

Sepsis is a life threatening health condition which occurs due to the severe and systemic inflammatory response to infection(Kingsley et al. 2020). The syndrome is initially characterized by dysregulated inflammatory, namely systemic inflammatory response syndrome (SIRS), progressing to severe sepsis and septic shock(Mayr, Yende, and Angus 2014). Sepsis is the most common cause of death for critically ill patients in non-coronary intensive care units(Angus et al. 2001). Necroptosis, leading to enormous release of inflammatory mediated and exacerbates inflammation, plays a vital role in the pathophysiology of systemic sepsis(Duprez et al. 2011; Hu et al. 2018). TNF-α, cytokine that plays a key role in sepsis, triggers necroptosis in response to tissue injury and inflammation(Xagorari, Roussos, and Papapetropoulos 2010). The most pertinent information regarding necroptosis was derived from studies of TNF-α induced necroptosis(Junying et al. 2019; Chen et al. 2014; Shan et al. 2018). Using a bioinformatics analysis and the drug repositioning strategy, we predicted a set of potential drug candidates for sepsis and validated the effectiveness of drug using both in vitro and in vivo models of TNF-α-induced necroptosis. Our experimental validation found that the candidate drug linifanib effectively inhibited necroptosis, SIRS and related tissue damage.

This research makes the following features. Firstly, we predict therapeutic drugs by analyzing the differences of gene expression in patients with sepsis. This bioinformatics prediction is an efficient drug screening method compared with traditional experiments. Secondly, by analyzing the pathophysiological molecular mechanism that plays an important role in sepsis, this study established an effective experimental verification system for the drug candidates predicted by bioinformatics. Thirdly, in this study, LINCS database containing FDA approved drug library was used, and then linifanib was repurposed to inhibit necroptosis, related inflammatory and SIRS. Since the safety evaluation of the drug has been completed, it is expected to be quickly transformed into clinical application.

Sepsis-induced acute lung injury remains the major cause of death in septic patients(Xu et al. 2019). It has been found that necroptosis is activated in the lung tissue of septic patients caused by New Coronavirus infection(Schroeder et al. 2019). RIPK1 kinase plays a crucial role in mediating necroptosis mechanisms upon the activation of TNFR1 by TNF-α(Junying et al. 2019). Upon necrosome formation, RIPK1 is activated and auto-phosphorylated, activating RIPK3 which binds to and phosphorylates the pseudokinase MLKL mediating necroptosis in disease pathogenesis(Alexei et al. 2019; Mifflin, Ofengeim, and Yuan 2020). Our study found that the expression of RIPK1, RIPK3 and MLKL in peripheral blood nucleated cells of sepsis patients was significantly higher than that of healthy controls. This provided further evidence that necroptosis plays a significant role in the pathophysiology of sepsis. Thus, RIPK1-dependent necroptosis may be a potential novel therapy target of sepsis.

Quantification of ADP-Glo kinase assays were performed with recombinant hRIPK1 in the presence of increasing concentrations of linifanib in our study. We found that linifanib inhibited the kinase activity of RIPK1 dose dependently, with an EC50 of 1182 nm, which was lower than that of the necroptosis inhibitor Nec-1 as a positive control. This indicates that linifanib may directly suppress RIPK1 kinase activity with excellent prospects for clinical application.

Necroptosis is characterized by marked swelling of cell, rupture of the plasma membrane and subsequent DAMPs released after a cellular disruption(Wang et al. 2015). TNF-α-induced necroptosis initiates downstream signaling cascades driving the production of a series of pro-inflammatory cytokines including IL-6(Kearney et al. 2015). Elevated blood concentrations of IL-6 triggered a detrimental cytokine storm(Chen et al. 2020) in sepsis were negatively correlated with the poor prognosis of patients(Chen et al. 2013). Endothelial cell necroptosis may contribute to lethality in both SIRS and sepsis patients(Zelic et al. 2018). Therefore, we adopted the SIRS mice model induced by tail vein injection of TNF-α to verify the role of candidate drugs from our previous bioinformatics analysis. Our study found that linifanib can effectively rescue the shock death and inhibit the overexpressed level of IL-6 mRNA in the damage lung tissue of SIRS mice. These data provide a solid basis for the clinical application of linifanib in the treatment of sepsis.

This study had two limitations. Firstly, despite recent research progress, our understanding of SIRS pathogenesis is still incomplete. More studies in the pathophysiological mechanism of sepsis and verification systems for therapeutic targets of sepsis are also needed. Secondly, whether linifanib can inhibit sepsis-related inflammatory response and reduce the mortality in patients need to be proved by clinical trials. However, linifanib is expensive before invalidation of the patents existed, thus limiting its current application in *investigator-initiated clinical trials* and translational research.

In conclusion, linifanib was demonstrated to inhibit RIPK1-dependent necroptosis and attenuate SIRS-induced acute lung injury. The present results provide further basis for the repurposing of linifanib to decrease sepsis mortality.

## Materials and Methods

### Drug prediction

#### Data collection

In order to predict new drugs for the treatment of sepsis, we performed bioinformatics analysis based on gene expression data of sepsis. The datasets associated with sepsis were retrieved and downloaded from the Gene Expression Omnibus (GEO, http://www.ncbi.nlm.nih.gov/geo/) database by setting the keyword as “sepsis”, organism as “Homo sapiens”, and study type as “expression profiling by array”. The datasets included in this study need to meet the following criteria: 1) the dataset should be human gene expression characteristics: and 2) gene expression data should be generated from sepsis patients and healthy subjects. Finally, the datasets GSE69528(Pankla et al. 2009), GSE46955(Shalova et al. 2015) and GSE54514(Parnell et al. 2013) were selected, as shown in Table 1. The samples selected from GSE69528 were blood samples from 29 patients with sepsis caused by B.pseudomallei and 28 healthy subjects. The samples selected from GSE46955 were monocytes from 8 patients with sepsis and 6 healthy subjects. The samples selected from GSE54514 were 31 blood samples from nonsurvivors of sepsis and 96 blood samples from survivors. These datasets were used to analyze differentially expressed genes between patients with sepsis and controls.

#### Identification of differentially expressed genes (DEGs)

Analysis of DEGs was conducted by using limma package (version 3.46.0) in R software (version x64 4.0.3). The downloaded gene expression data has been log2 transformed, and we used R language to process the data. DEGs were identified with a level of *P* value < 0.05. And the heatmaps of the top 100 DEGs was produced by pheatmap package (version 1.0.12) and R software.

#### Identification of signaling pathway

In order to identify the key pathogenic signaling pathways associated with sepsis, we searched the literature with the keyword “sepsis” in PubMed, and identified the signaling pathways closely related to the pathogenesis of sepsis from the literature.

#### Perturbation-response signatures of drugs

The Library of Integrated Network-Based Cellular Signatures (LINCS) contains the characteristics of drug functional genomics and drug metabolomics of individual cells. Human cell lines were treated with small molecular compounds, and then the multi-level cell responses before and after cell treatment were compared(Keenan et al. 2018).

There is a high correlation between human gene expression, and it is possible to calculate the expression levels of different genes. The L1000 library in LINCS makes use of the correlation between gene expression, it mainly uses small molecular compounds to treat human cell lines, then 978 genes were identified as genome-wide marker genes in 384 well plate based on large-scale statistical analysis. The expression of other genes can be calculated by measuring the expression of marker genes. The experimental results show that the changes of nearly 1000 genes adopted by L1000 can represent about 80% of the gene change information of human(Subramanian et al. 2017).

The gene set of drug perturbation experimental data in LINCS can be divided into three parts: (1) Landmark space. It contains 978 landmark genes, which is based on the probe fluorescence intensity data measured by the experiment, scaled and standardized after calculation with the control group. (2) All inferred genes. It contains 978 landmark genes and 11350 genes whose expression data are inferred. There are 12328 genes. (3) Best inferred genes. It contains 978 landmark genes and 9196 high fidelity inferred genes from 11350 inferred genes. There are 10174 genes in this gene set, which is also the gene set used in drug prediction.

The LINCS data used in this study is the level 5 data “gse70138_broad_lincs_level5_compz_n1180 50×12328. Gctx. GZ” of GSE70138. For each experimental group, calculate the gene differential expression value under each perturbation experiment of small molecular compounds, the differential expression values are arranged from large to small. The perturbation data of each small molecular compound includes the gene expression data of different cell lines after different doses and different times of treatment.

#### Kolmogorov–Smirnov test

In statistics, Kolmogorov-Smirnov test(KS test) is a nonparametric test, which is based on cumulative distribution function to test whether an empirical distribution conforms to a theoretical distribution or compare whether there is significant difference between two empirical distributions (Hollander, Wolfe, and Chicken 2013; Lamb et al. 2006).

*X_1_, X_2_,…, X_m_, Y_1_, Y_2_,…, Y_n_* are two independent random samples, and the distribution functions are *F_m_*(*t*) and *G_n_*(*t*), they can be defined as:

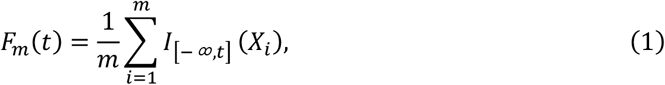

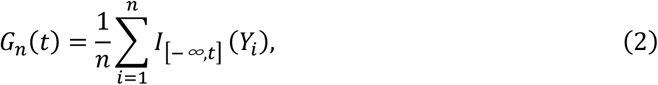

Where m and n denote the number of samples in the two groups, t denotes any real number, *I*_[−∞, *t*]_(*X_i_*) and *I*_[−∞, *t*]_(*Y_i_*) are defined as:

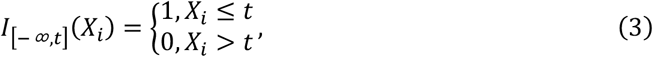

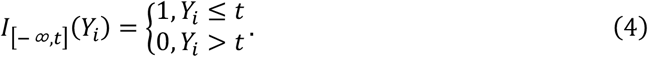

And the statistic of KS test is defined as:

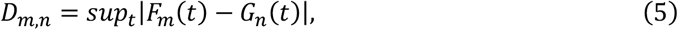

Where *sup_t_* is the supremum of distance.

The statistic of Kolmogorov Smirnov test quantifies the distance between the empirical distribution functions of two groups of samples. null hypothesis H0: the two data distributions are consistent or the data conform to the theoretical distribution. When the actual observed value *D_m,n_* is greater than the critical value, H0 is rejected, otherwise, H0 is accepted.

#### Calculation of the therapeutic scores

We use the differential expression information of genes in sepsis-related pathways to construct disease signatures, and use the differential expression information of genes under the action of drugs to construct drug signatures. The differential expression values of genes in sepsis-related pathways came from log-fold change (logFC) after differential expression analysis of sepsis-related datasets, and the gene differential expression information under the action of drugs comes from L1000 library. Based on the statistic of KS test, these disease signatures and drug signatures are used to design a pattern matching method to calculate the treatment scores of different drugs on sepsis.

For sepsis-related signaling pathway C and drug-induced gene expression profile D, the treatment scores of up-regulated gene set and down-regulated gene set of pathway C relative to drug-induced gene expression profile D were calculated respectively.

Firstly, take the intersection of up-regulated genes or down-regulated genes in sepsis-related pathways and genes under drug perturbation, record this gene set as H. The genes induced by drugs were arranged according to the differential expression value from large to small, and the genes in H were arranged from large to small according to this differential expression value. Using the statistic of KS test, a and b are defined as follows:

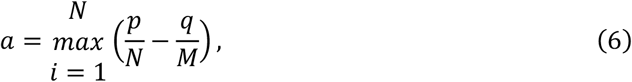

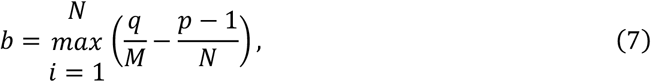

Where N is the total number of genes in the H gene set, and M is the total number of genes whose expression value changes under drug perturbation, and P is the position of current gene g in gene set H, and Q is the position of gene g in the gene list under drug perturbation.

Therefore, the treatment score of up-regulated gene set or down-regulated gene set in pathway C is calculated as

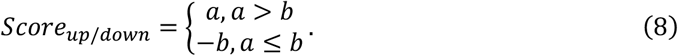

And the treatment score of drug for sepsis-related signaling pathway C is calculated as

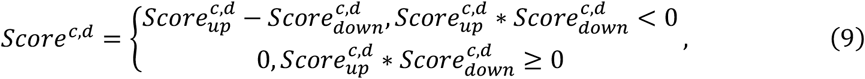

Where c denotes the sepsis-related signaling pathway and d denotes the drug.

There are four possible effects of drugs on a sepsis-related signaling pathway.

In the first case, the change of gene expression under drug perturbation is opposite to that under sepsis, so this drug has the potential to treat sepsis, in this case, *Score^c,d^* < 0. In the second case, for the genes whose expression value decreases in sepsis, under drug perturbation, the expression value of some genes increases and the expression value of others decreases. Then the use of the drug in sepsis will cause the expression of some genes to return to normal, but the expression of other genes will continue to decrease. In the third case, for genes with increased expression value in sepsis, under drug perturbation, the expression value of some genes increases and the expression value of others decreases. Then the use of the drug in sepsis will cause the expression of some genes to return to normal, but the expression of other genes will continue to increase. In the fourth case, the gene whose expression value increases or decreases in sepsis also increases or decreases under the action of the drug, indicating that the effect of the drug and sepsis on gene expression changes is consistent, and the drug may aggravate the severity of sepsis, in this case, *Score^c,d^* > 0.

In the first and fourth cases, 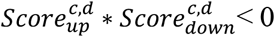. After screening the drugs that meet these two conditions, the drugs meeting the first condition were further screened according to *Score^c,d^* < 0. The greater the absolute value of *Score^c,d^*, the more obvious the effect of the drug.

After calculating the drug score for each pathway, further processing was performed: Scores obtained by treating the same cell line with a drug at different times and doses, Only the results with the largest absolute score are retained, scores obtained by treating different cell lines with a drug, the mean value of the score was calculated and its absolute value was used as the treatment score of the drug.

For each sepsis-related dataset, the average score of each drug in all pathways was calculated. Only results with score < 0 are retained, and sorted according to absolute value of the score from large to small, including only the drugs with potential therapeutic effect on sepsis. Finally, take the overlapping parts of the Top60 drug list of three datasets to determine the final candidate drugs.

### Experimental verification

Biological reagents. The reagents used were as follows: Flag-mTNF-α was a gift from professor Daichao Xu (Interdisciplinary Research Center on Biology and Chemistry, Shanghai). We used the following reagents: recombinant murine / human TNF-α (Sino, 50349-MNAE) and (Peprotech, 300-01A-100); Linifanib (Selleck Chemicals, S1003, CAS# 796967-16-3); Pierce protease inhibitor tablets (Thermo Fisher Scientific, A32965); z-VAD-fmk (Selleck Chemicals, S7023); Smac mimetic (SM-164) (Beyotime Biotechnology, SC0114-10mM); Necrostatin-1 (Selleck Chemicals, S8037); CellTiter-Glo Luminescent Cell Viability Assay (Promega, G7573); FDA-approved chemical library (Selleck Chemicals, L1300); ANTI-FLAG® M2 Affinity Gel (Sigma, A2220); Protein A/G ultra link resin (Thermo Fisher Scientific, 53133); Pierce Protease Inhibitor Tablets (Thermo Fisher Scientific, A32965).

The antibodies used for Western were obtained from commercial sources: anti-RIPK1 (BD Biosciences, 610459 and Cell Signaling Technology, 3493S); anti-mouse phosphoS166-RIPK1 (Cell Signaling Technology, Cat#31122S); anti-mouse MLKL (Proteintech, 66675-1-Ig); anti-mouse-phosphoS345-MLKL (Abcam, ab196436), anti-Tubulin (Proteintech, 66031-1-Ig); anti-A20 (Cell Signaling Technology, Cat#5630); anti-TNFR1 (Proteintech, 21574-1-AP); anti-TAK1(Proteintech, 12330-2-AP);anti-human phospho-RIPK1 (Cell Signaling Technology, Cat#65746), anti-human MLKL (Abcam, Cat#ab183770), anti-human phospho-MLKL (Abcam, Cat#ab187091).

#### Cells and cell culture

Mouse embryonic fibroblasts (MEFs) and FADD-deficient Jurkat cell line were kindly provided by Prof. Daichao Xu of Interdisciplinary Research Center on Biology and Chemistry, CAS, Shanghai, China. MEFs were cultured in RPMI 1640 medium (Hyclone). Medium were additionally supplemented with 10% FBS (Zhejiang Tianhang Biotechnology), MEM (Hyclone), NEAA (Hyclone), and antibiotics (100 U/ml penicillin and 100 mg/ml streptomycin) (Gibco). All of cell lines were cultured in a humidified 5% CO2 atmosphere at 37 °C.

#### Necroptosis induction and cell viability analysis

For FADD - deficient Jurkat cells, necroptosis was induced by human TNF-α (50 ng/ml) for 24 hr. For MEFs cells, necroptosis was induced by pretreatment with Smac164 (200 nM) and z-VAD-fmk (20 μM) for 30 min and followed by mTNF-α (25 ng/ml). The compounds were incubated with the cells at the indicated concentrations for 24 hr. Cell viability was then examined by using the CellTiter-Glo Luminescent Cell Viability Assay kit (Promega). Luminescence or absorbance was recorded with a BioTek 312e microplate reader (BioTek Instruments, Winooski, VT).

#### Animal experiments

All animal care and experimental procedures complied with the National Institutes of Health guidelines (NIH publications Nos. 80–23, revised 1996) and under the approval of the Ethical Committee of the Three Gorges University Laboratory Animal Center. For TNF-induced SIRS model, male C57BL/6 J mice (8–10 weeks old) were purchased from Beijing Vital River Laboratory Animal Technology Co., Ltd. (Beijing, China). Mice were raised in a SPF facility with a pathogen-free environment (23 ± 2°C and 55 ± 5% humidity) of 12:12-hour light/dark cycle at the Three Gorges University Laboratory Animal Center.

The animal protocols were approved by the Standing Animal Care Committee at China Three Gorges University. Compounds were suspended in distilled water containing 0.5% carboxymethyl cellulose sodium. In drug treatment groups, mice were given linifanib by p.o. gavage 30min before TNF injection. Mouse TNF-α (mTNF-α) were diluted in endotoxin-free PBS and injected i.v. (5ug/mice) in a volume of 200μl. Rectal body temperature was recorded with an electric thermometer. Lung tissues were collected at indicated times after sacrificing the mice. At the end of the experiment, mice were sacrificed with an overdose of anaesthetic (i.v. injection of 1.2% avertin solution; 0.2 ml per 10 g).

#### RNA extraction, reverse transcription, and real-time PCR

The mRNA levels of lung tissues from SIRS model mice were analyzed at 6h after injection of TNFα. Total lung RNA was extracted with the FastPure Cell/Tissue Total RNA Isolation Kit V2 (RC112-01, Vazyme) and reverse transcribed into cDNA using the HiScript III RT SuperMix (R323-01, Vazyme). Quantitative real-time PCR was performed using ChamQ SYBR qPCR Master Mix (Q311-02, Vazyme) and analyzed with CFX Manager software of Bio-rad CFX384. Relative gene expression was normalized to 18s and determined using the -ΔΔCT method. The primer sequences were as following: for 18s, forward: 5’-AGTCCCTGCCCTTTGTACACA-3’ and reverse 5’ - CGATCCGAGGGCCTCACTA-3’; for IL-6, forward: 5’ - TACCACTCCCAACAGACCTG-3’ and reverse 5’-GGTACTCCAGAAGACCAGAGG - 3’.

#### Western blotting analysis

Cells were lysed in 1% NP-40 buffer for total lysis (1% Nonidet P-40, 50 mM Tris Base (pH 7.5), 150 mM NaCl, 1 mM PMSF, 1 mM NaF/Na3VO4, Pierce Protease Inhibitor Tablets). All cell lysis buffers were supplemented with 1% NP-40 buffer to the same concentration and were then added with 5x loading buffer (10% SDS, 40% glycerol, 25% 1M 336 Tris6 -HCl (pH 6.8), 0.005% bromophenol blue, 25% β-Mercaptoethanol) with denaturation at 95°C for 5 min. Cell lysates were analysed by SDS-PAGE gels with running buffer and subsequently transferred onto immobilon-NC transfer membrane (NC membrane, HATF00010, Millipore) or 0.45 μM polyvinylidene difluoride membrane (PVDF membrane, MXHVWP124, Millipore). Membranes were blocked in 5% skim milk in Tris-buffered saline (TBS) containing 0.1% Tween 20 before overnight incubation with specific primary antibodies at 4°C. All listed primary antibodies were used at 1:1000. Membranes were then washed and incubated with appropriate HRP-conjugated secondary antibodies (1031-05 and 4050-05, SouthernBiotech), developed immunoreactivity (G2014-50ML, Servicebio), and imaged using the Tanon-4800 345 (Tanon Science & Technology Co., Ltd.).

#### Immunoprecipitation (IP)

For immunoprecipitation (IP) of complex I, MEFs were lysed by 1% NP-40 buffer (1% Nonidet P-40, 50 mM Tris Base (pH 7.5), 150 mM NaCl, 1 mM PMSF, 1 mM NaF/Na3VO4 and Pierce Protease Inhibitor). Complex I was purified by IP using ANTI-FLAG® M2 Affinity Gel of 20ul. The IP protein was rotated overnight in 4°C. The beads were washed three times with ice-cold 4% NP-40 buffer and eluted by directly adding 2x loading buffer (4% SDS, 20% glycerol, 10% 1 M Tris-HCl (pH 6.8), 0.005% bromophenol blue, 10% b-Mercaptoethanol) in 95°C for 5 min, then analyzed by immunoblotting (antibodies used as indicated).

#### In vitro kinase assays

We followed the protocol of RIPK1 Kinase Enzyme System (Promega, Cat#VA7591) and ADP-Glo™ Kinase Assay (Promega, Cat# V6930) to detect the inhibition of RIPK1 kinase activity by linifanib.

#### Data and statistical analysis

The data and statistical analysis comply with the recommendations of the British Journal of Pharmacology on experimental design and analysis in pharmacology. Results are presented as means ± SEM. Student’s t-test and one-way ANOVA were used for comparison among the different groups. The log-rank (Mantel-Cox) test was performed for survival curve analysis using GraphPad Prism8.0. P < 0.05 was considered statistically significant.

## Acknowledgments

Thanks to all those who maintain excellent databases and to all experimentalists who enabled this work by making their data publicly available. We would like to thank Dr. Hongbing Zhang of Peking Union Medical College for his mentoring and guidance in the field of drug repurposing. This work was supported by the National Natural Science Foundation of China (62072353). This work was supported by grants from Health Commission of Hubei Province Scientific Research Project (WJ2021M257), Local Development Project of Science and Technology guided by the Central Commission (ZYYD2020000202), Project of Hubei Province Clinical Medical Research Center for Rare Diseases of Nervous System, Hubei Province Outstanding Medical Academic Leader Program (EWT201947), Yichang Famous Doctor Studio, Yichang Training Talents of Innovation Entrepreneurship and Excellence-creating project (JY201701) and Project of Yichang City Medical and Health Research (A22-2-031).

